# Revision of the genus *Dichaetophora* Duda (Diptera: Drosophilidae), part I: DNA barcoding and molecular phylogenetic reconstruction

**DOI:** 10.1101/2021.05.28.446102

**Authors:** Takehiro K. Katoh, Ji-Min Chen, Jin-Hua Yang, Guang Zhang, Lu Wang, Awit Suwito, Masanori J. Toda, Ya-Ping Zhang, Jian-Jun Gao

## Abstract

The genus *Dichaetophora* is currently comprised of 67 formally described species assigned into five species groups, i.e., *agbo*, *tenuicauda*, *acutissima*, *sinensis* and *trilobita*. In the present study, we challenged to delimit species from a huge amount of samples of *Dichaetophora* and allied taxa (the genus *Mulgravea* and the subgenus *Dudaica* of *Drosophila*) collected from a wide range of the Oriental and east Palearctic regions. We first sorted all specimens into morpho-species, which were tentatively classified into the species groups of *Dichaetophora*, *Mulgravea* and *Dudaica* based on the morphological diagnoses for these supraspecific taxa. Then, we selected representative specimen(s) for each morpho-species and subjected them to barcoding of *COI* (the cytochrome *c* oxidase subunit I gene) sequences. The applied ABGD (automatic barcode gap discovery) algorithm estimated a total of 222 MOTUs (molecular operational taxonomic units). Although most of the morpho-species were recognized as individual MOTUs, some others were divided into multiple MOTUs or combined into a single MOTU, suggesting the existence of cryptic sibling species, subspecies or intraspecific genetic varieties. Out of the 222 MOTUs, 88 representing the supraspecific taxa of *Dichaetophora*, *Mulgravea* and *Dudaica* were selected, along with 33 species from major genera and subgenera of *Drosophila* in the tribe Drosophilini, as in-group (four species from the tribe Colocasiomyini as out-group) for phylogenetic reconstruction. We analyzed a dataset of concatenated nucleotide sequences of 12 nuclear gene markers by the maximum likelihood and Bayesian inference methods. As a result, the three focal taxa (i.e., *Dichaetophora*, *Mulgravea* and *Dudaica*) formed a clade, which we called the “pan-*Dichaetophora*”. Within this large clade, the *agbo*, *tenuicauda*, *sinensis* and *trilobita* groups of *Dichaetophora*, *Mulgravea* and *Dudaica* were recovered as monophyletic groups, but *Dichaetophora* and its *acutissima* group were regarded as paraphyletic. In addition, two clusters were recognized among ungrouped MOTUs of *Dichaetophora*. Thus, the present study has uncovered some problems in the taxonomy of the pan-*Dichaetophora*. Solving such problems will be challenged in Part II of this serial work.

## Introduction

Duda (1940) established *Dichaetophora* as a subgenus in the genus *Drosophila* Fallén (Drosophilidae: Drosophilinae) with *Drosophila aberrans* Lamb, 1914 from the Seychelles as the type species. After that, a number of species were added to this subgenus from Africa (3 spp.; Burla, 1954; Graber, 1957) and East Asia (6 spp.; Lee, 1964; Okada, 1964, 1966, 1968; Kang *et al.*, 1967). These East Asian species were, however, later transferred to the genus *Nesiodrosophila* Wheeler & Takada (Okada, 1976, 1977, 1984a), which was established with *Nesiodrosophila lindae* Wheeler & Takada, 1964 as the type species by Wheeler & Takada (1964). Then, a large number of *Nesiodrosophila* species were found from the Old World: 14 spp. from the Oriental region (Lin & Ting, 1971; Okada, 1984a, 1988; Gupta & De, 1996), 3 spp. from the Palearctic region (Nishiharu, 1981; Toda, 1989), 15 spp. from the Australasian region (Bock, 1982; Okada, 1984a; Toda *et al.*, 1987), and 1 sp. from the Afrotropical region (Okada, 1984a). In addition, another related taxonomic group, i.e., the *Drosophila tenuicauda* species group, was recognized within the subgenus *Lordiphosa* Basden of *Drosophila* (Toda, 1983; Okada, 1984b; Hu *et al.*, 1999; Katoh *et al.*, 2000): Hu & Toda (2001) inferred the sister relationship between the *tenuicauda* group and *Nesiodrosophila* from a phylogenetic analysis based on 68 morphological characters. Grimaldi (1990) elevated *Dichaetophora* and *Lordiphosa*, along with the subgenera *Hirtodrosophila* Duda and *Scaptodrosophila* Duda of *Drosophila*, to the generic rank, based on the results of a family-wide cladistic analysis on 217 adult morphological characters.

Taking into account these possible relationships proposed in the previous studies, Hu & Toda (2002) conducted a morphological cladistic analysis focusing on *Dichaetophora*, *Nesiodrosophila* and the *Lordiphosa tenuicauda* group. Based on the result that these three taxa formed a monophyletic group, all the species so far assigned to them were merged into the revised genus *Dichaetophora*: i.e., *Nesiodrosophila* was synonymized with *Dichaetophora*. And, within the revised *Dichaetophora*, three species groups were newly proposed: the *agbo* group comprised the four Afrotropical *Dichaetophora* species and all the species so far assigned to *Nesiodrosophila*, and the previous *Lo. tenuicauda* group was split into the *tenuicauda* and *acutissima* groups. Since then, two more species groups were added to *Dichaetophora*: the *sinensis* group comprised of four Chinese species (Hu & Toda, 2005) and the *trilobita* group of six Oriental species (Yang *et al.*, 2017).

The current genus *Dichaetophora* includes a total of 67 formally described species: 43 spp. of the *agbo* group, 10 spp. of the *tenuicauda* group, 4 spp. of the *acutissima* group, 4 spp. of the *sinensis* group, and 6 spp. of the *trilobita* group; 5 spp. distributed in the Palearctic region (East Asia), 6 spp. in the Palearctic (East Asia) and Oriental regions, 35 spp. in the Oriental region, 15 spp. in the Australasian region, 5 spp. in the Afrotropical region, and 1 sp. in the Palearctic (East Asia), Oriental and Australasian regions (DrosWLD-Species: https://bioinfo.museum.hokudai.ac.jp/db/index.php; TaxoDros: http://www.taxodros.uzh.ch/). Our intensive surveys of drosophilid faunas in the Oriental region during the past two decades uncovered very high species diversity of *Dichaetophora* from previously less explored microhabitats: an unexpectedly large number of putatively new species (*ca*. 150) were recognized among specimens collected by net sweeping mostly from herbaceous stands or forest floor, occasionally from tree-trunks, flowers, fallen fruits and fungi, or by light traps. This finding of the great species diversity prompted us to a revisional work on the systematics of *Dichaetophora*.

The phylogeny of *Dichaetophora* has been less explored. Grimaldi (1990) included two species, *Dr*. *aberrans* and *Nesiodrosophila rotundicornis* (Okada, 1966), of the current *Dichaetophora* in his morphological cladistic analysis with an extensive taxon sampling of most genera and subgenera of the family Drosophilidae. However, the two species were placed in different lineages distant from each other in the resulting tree. When Hu & Toda (2002) redefined the genus *Dichaetophora*, they suggested its relationships with the genera *Jeannelopsis* Séguy, *Sphaerogastrella* Duda, *Mulgravea* Bock and *Liodrosophila* Duda because an important diagnostic character “the oviscapt with apical ovisensillum robust and the largest, distinguishable from the others” for *Dichaetophora* is shared as a synapomorphy (ap. 213) of Grimaldi (1990) by these genera. This character is shared by the subgenus *Dudaica* Strand of *Drosophila* as well (Katoh *et al.*, 2018). On the other hand, Hu & Toda (2005) suggested the sister relationship between *Hirtodrosophila* and the monophyletic *Dichaetophora* comprised of the *agbo*, *tenuicauda*, *acutissima* and *sinensis* groups. The close relationship between *Dichaetophora* (the *tenuicauda* and *acutissima* groups) and *Hirtodrosophila* was suggested in molecular phylogenetic analyses by Katoh *et al.* (2000) and Russo *et al.* (2013) as well. Yassin (2013) constructed a family-wide Bayesian phylogenetic tree, based on a multi-locus (seven nuclear and one mitochondrial genes) dataset of DNA sequences from 190 species of 33 drosophilid genera, and inferred that *Dichaetophora* (the *agbo*, *tenuicauda* and *acutissima* groups) formed a cluster with the genera *Hirtodrosophila*, *Mycodrosophila* Oldenberg, *Zygothrica* Wiedemann, *Dettopsomyia* Lamb and *Jeannelopsis*. However, the statistical supports (Bootstrap, Bremer and/or Posterior Probability values) for these relationships were all low. Thus, the phylogenetic position of *Dichaetophora* and the relationship within this genus are still to be investigated, especially by means of molecular phylogenetic methods.

In the present study, we first try to delimit species of *Dichaetophora* and its relatives (the genus *Mulgravea* and the subgenus *Dudaica* of *Drosophila*) based on morphological and DNA barcode data, examining a huge amount of specimens of known and putatively new species of these focal taxa collected by our surveys for ten-odd years in the Oriental and East Asian regions. We then conduct a multi-locus molecular phylogenetic analysis with taxon sampling expanded from the three focal taxa to possibly allied genera/subgenera such as those mentioned above and species representing some major lineages of the subfamily Drosophilinae. The phylogeny to be reconstructed in this study will provide a framework for revising the taxonomy of *Dichaetophora* and allied taxa in a subsequent study of this serial work and baseline knowledge for future evolutionary studies on this speciose group of drosophilids adapted to particular microhabitats.

## Materials and methods

### Fly samples and species delimitation

We selected the genera *Dichaetophora* and *Mulgravea* and subgenus *Dudaica* of *Drosophila* as focal taxa from our samples of drosophilid flies collected from East and Southeast Asia (China, Japan, Vietnam, Myanmar, Malaysia and Indonesia) and Australia, mostly by net sweeping over herbaceous stands along forest edges or on the forest floor, sometimes by aspirating from flowers or by using light traps set in the tree canopy. Specimens were preserved in either ethanol (70% or 100% for morphological or molecular study, respectively) or Kahle’s solution (to maintain the pigmentation of specimens for a long time in the laboratory; only for specimens collected from Myanmar and Australia) before this study.

The selected specimens were first sorted into morpho-species. A few representative specimens of each morpho-species were dissected by detaching the genitalia and/or some other body parts (e.g., head, mouthparts, legs, etc.) from the main body, and the detached organs were treated and examined in the same method as that described by Shi *et al.* (2019). And, putatively new species were tentatively classified into the five species groups of *Dichaetophora*, *Mulgravea* or *Dudaica*, based on their diagnostic, morphological characters (Hu & Toda, 2002, 2005 for the *agbo*, *tenuicauda*, *acutissima* and *sinensis* groups of *Dichaetophora*; Yang *et al.*, 2017 for the *trilobita* group of *Dichaetophora*; Bock, 1982 for *Mulgravea*; Katoh *et al.*, 2018 for *Dudaica*). In this classification, many morpho-species completely conformed to the diagnosis of any group (i.e., having all the diagnostic characters), but some others only partially, and those of which morphology were inconsistent with any diagnosis were treated as ungrouped.

Some of the specimens thus identified to morpho-species were selected and subjected to DNA barcode sequencing, considering the total number, gender and geographical origins of the available specimens for each morpho-species (Table S1). We used the same methods as in Yang *et al.* (2017) for fly tissue sampling, DNA extraction (using TIANamp^®^ Genome DNA Kit) and PCR [using TIANGEN^®^ *Taq* DNA polymerase and Folmer *et al.*’s (1994) primer pair LCO1490/HCO2198]. The PCR products were subjected to sequencing in the TSINGKE Biological Technology (http://www.tsingke.net) with an ABI 3700 sequencer, with trace files edited subsequently in the SeqMan module of the DNAStar package version 7.1.0 (DNAStar Inc., Madison, WI).

Newly determined barcodes were aligned with 78 *COI* sequences downloaded from GenBank (Table S1) in MEGA 7 (Kumar *et al.*. 2016) with the ClustalW method. A neighbor-joining (NJ) tree was then built with the resulting sequence alignment in MEGA 7 (options: model = *p-*distance, variance estimation method = bootstrap method of 1,000 replicates, gaps treatment = pair-wise deletion). Recognition of MOTUs (molecular operational taxonomic unit) was then conducted using the ABGD (automatic barcode gap discovery) algorithm (Puillandre *et al.*, 2012) through the web interface (https://bioinfo.mnhn.fr/abi/public/abgd/abgdweb.html), with the “simple distance” (i.e., *p-*distance) option under default settings: Pmin = 0.001, Pmax = 0.1, Steps = 10, X (a proxy for minimum gap width) = 1.5, Nb bins (for distance distribution) = 20.

### Taxon sampling for phylogenetic reconstruction

We expanded the taxon sampling from the three focal taxa to some previously suggested relatives, such as *Hirtodrosophila*, *Mycodrosophila*, *Zygothrica*, *Dettopsomyia* and *Liodrosophila*, and also some major lineages, such as the subgenera *Drosophila*, *Siphlodora* Patterson & Mainland and *Sophophora* Sturtevant of the genus *Drosophila* and Hawaiian drosophilids (*Idiomyia* Grimshaw and *Scaptomyza* Hardy), of the tribe Drosophilini Okada (*sensu* Yassin, 2013) as in-group taxa. For the three focal taxa, not all but a part of MOTUs recognized in the ABGD analysis of *COI* barcodes were applied to the phylogenetic reconstruction by selecting representatives from each genus, subgenus, species group and/or cluster recognized in the barcoding tree. In total, 88 MOTUs (including 25 known species) from the three focal taxa and 33 species from other in-group genera/subgenera were sampled for the phylogenetic analysis (Table S2). As out-group taxa, four species of *Chymomyza* Czerny, *Colocasiomyia* de Meijere, *Impatiophila* Fu & Gao and *Scaptodrosophila* were selected from the tribe Colocasiomyini Okada (*sensu* Yassin, 2013) (Table S2).

### Genetic markers and DNA sequencing

We adopted 12 nuclear gene markers: the D7 region of the *28S rRNA* locus (Friedrich & Tautz, 1997) was used as a practicable marker, and 11 single-copy, orthologous, protein-coding gene (PCG) markers were set up by referring to the alignments of our unpublished transcriptome sequences of ten drosophilid species (three of *Dichaetophora* and seven of allied genera). The PCG markers were evaluated on the basis of sequence variation within the coding part of the target regions of the alignments.

We then designed PCR/sequencing primer pairs for each of these markers (Table 1) and chose optimal reaction condition for each pair by evaluating their performance with DNA samples of eight *Dichaetophora* species, *acutissima* (Okada, 1956), *pseudocyanea* (Hu & Toda, 1999), *tenuicauda* (Okada, 1956), *facilis* (Lin & Ting, 1971), *ogasawarensis* (Toda, 1987), *lindae*, *neocirricauda* (Gupta & De, 1996) and *trilobita* Yang & Gao, 2017, covering all the five species groups of this genus. Then, the target regions of the selected nuclear markers were amplified and sequenced for the in- and out-group taxa with the designed primer pairs (with the same procedures described in the DNA barcoding section, in the TSINGKE Biological Technology), and the resulting trace files were edited with SeqMan or MEGA7. The protein-coding part(s) of the newly collected sequences of each locus were aligned with some homologous sequences available from GenBank using the ClustalW method, and then concatenated for all loci, also in MEGA 7.

**Table 1.**
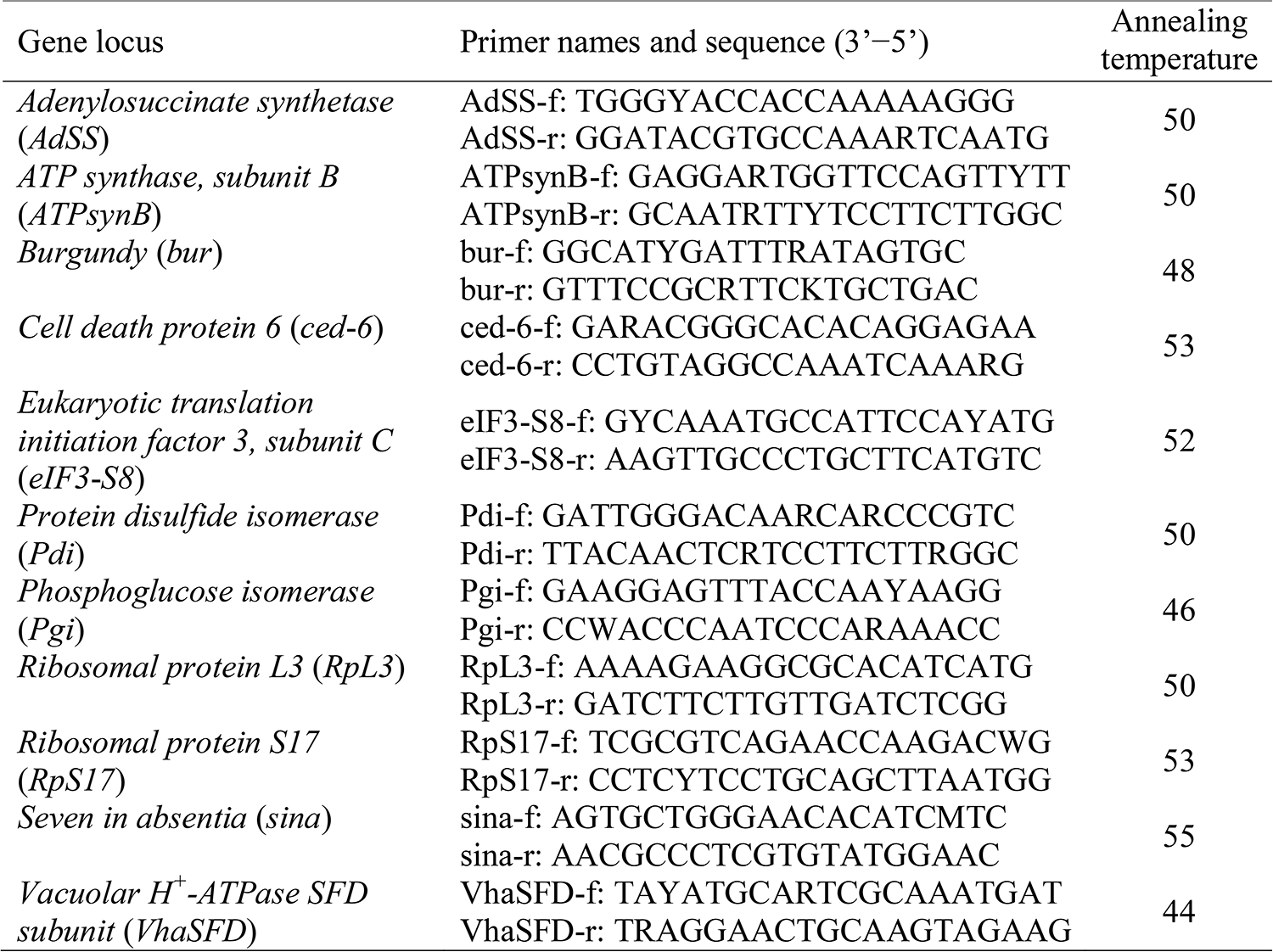
PCR/sequencing primer pairs designed for nuclear markers in the present study.

### Data partitioning and multi-locus phylogenetic reconstruction

We used the program PartitionFinder 2 (Lanfear *et al.*, 2016) to search optimal partitioning scheme for the alignment of the concatenated sequences and select the best fit nucleotide substation model for each suggested partition. The search was conducted with the greedy algorithm (Lanfere *et al.*, 2012) under the BIC (Bayesian Information Criterion; Schwarz, 1978), with data blocks defined in light of gene locus and each codon position for the PCG sequences. The model was set as “models=all” for the ML, but “model=mrbayes” for the BI analyses. The ML tree was constructed using RAxML 1.0.1 (Kozlov *et al.*, 2019), with node confidences evaluated through 1,000 non-parametric bootstrap replicates. The BI analysis was performed in MrBayes 3.2.7 (Ronquist *et al.*, 2012) through two independent runs each including four MCMC chains, with chains sampled every 100 generations. The convergence of runs was evaluated in Tracer V1.6 (Rambaut *et al.*, 2014) after discarding the initial 25% samples as burn-in.

## Results

### COI *barcoding*

We determined *COI* sequences for 1,013 specimens of the three focal taxa: 1,004 of *Dichaetophora*, 5 of *Mulgravea*, and 4 of *Dudaica* (Table S1). The NJ barcode tree built with the alignment of these barcodes and 57 GenBank *COI* sequences (23 of the *Di. trilobita* group and 34 of *Dudaica*; Table S1) is shown in Fig. 1 (partially compressed) and Fig. S1 (original). In the ABGD analysis with all but one (#00676 with only 212 nucleotide sites determined) of the *COI* sequences, a total of 177 to 311 MOTUs were recognized when the prior maximum intraspecific divergence (*P*) was changed from 0.001000 to 0.021544 (Table 2). We selected *P* = 0.004642 as an optimal, where the numbers of MOTUs identified in the initial and recursive partitions were 222 and 226, respectively. We adopted the former estimate (222) as a hypothetical number of MOTUs. Among them, most of morpho-species (including all the studied, known species) were recognized as individual MOTUs, but some others were divided into multiple MOTUs or two morpho-species (*Di.* sp.DLS3a and *Di.* sp.DLS3b) were combined into a single MOTU (Fig. 1, Fig. S1).

**Table 2.** The numbers of MOTUs estimated at various values of the prior intraspecific divergence (*P*) in the ABGD analysis.

| Prior intraspecific divergence ($P$) | Number of MOTUs | |
| --- | --- | --- |
|  | Initial partition | Recursive partition |
| 0.001000 | 309 | 311 |
| 0.001668 | 309 | 310 |
| 0.002783 | 309 | 310 |
| 0.004642 | 222 | 226 |
| 0.007743 | 222 | 224 |
| 0.012915 | 173 | 179 |
| 0.021544 | 173 | 177 |

**Figure 1.**
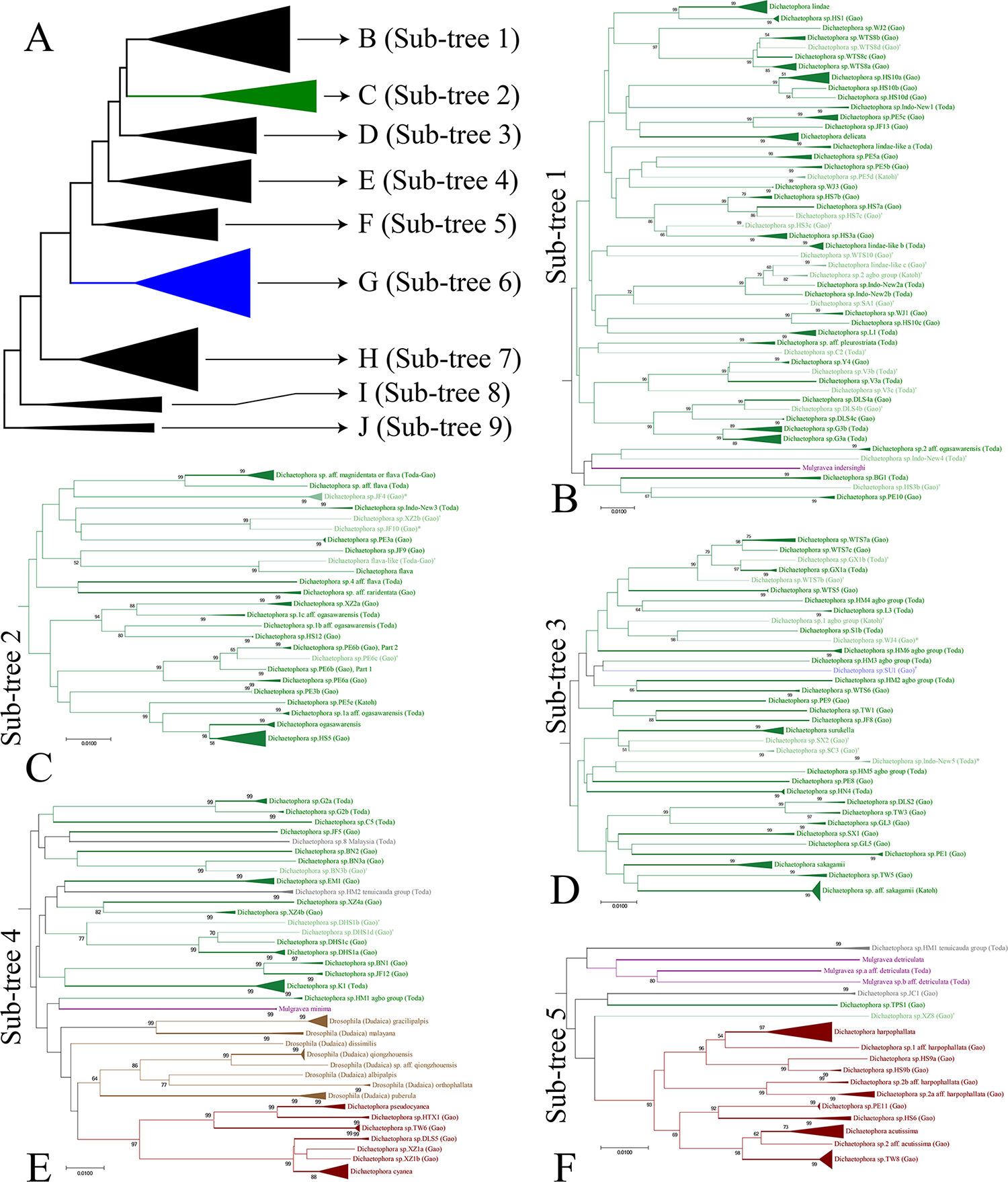
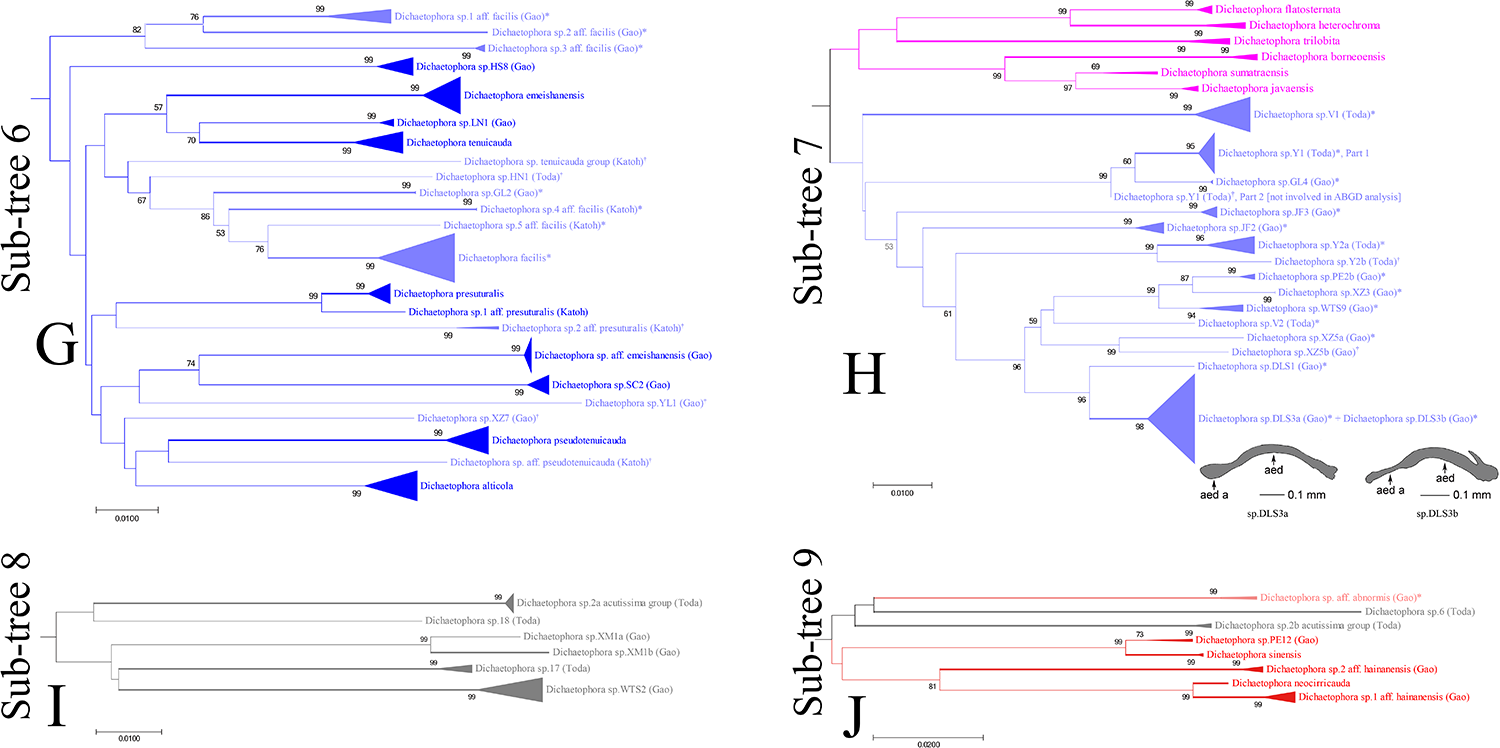
Neighbor-joining tree (within-MOTU branches compressed) built with 1,070 *COI* sequences of the genera *Dichaetophora* and *Mulgravea* and the subgenus *Dudaica* of *Drosophila*. The skeleton of the tree is shown in panel A, and sub-trees shown individually in B–J. Voucher specimen number (or GenBank accession number) is shown for each sequence. MOTUs inferred from the ABGD analysis is shown, with indication of supraspecific classification in different colors: the *agbo* (green), *acutissima* (brown), *tenuicauda* (blue), *trilobita* (pink), *sinensis* (red) species groups and ungrouped species (gray) of *Dichaetophora*, *Mulgravea* (purple), and *Dudaica* (light brown). MOTUs only partially concordant with the diagnosis are asterisked and those having female specimens only are indicated with dagger (†), and both are shown in pale color. MOTUs subjected to the phylogenetic reconstruction are shown with thick branch(es). Bootstrap percentages more than 50% are shown beside nodes.

### Phylogenetic reconstruction

DNA sequences of the 12 nuclear gene markers were newly determined for the 89 specimens (representing 88 MOTUs) selected from the three focal taxa (82 of *Dichaetophora*, three of *Dudaica*, and four of *Mulgravea*), 13 in-group and three out-group species (Table S2). The alignment of the data sets of these newly determined sequences and those from GenBank for 21 other species (20 in-group and 1 out-group spp.; Table S2) spans 4,571 nucleotide sites, among which 2,005 are variable and 1,710 are parsimony informative. The optimal partitioning strategy for the concatenated data set is shown in Table 3, together with the selected nucleotide substitution model for each partition.

**Table 3.**
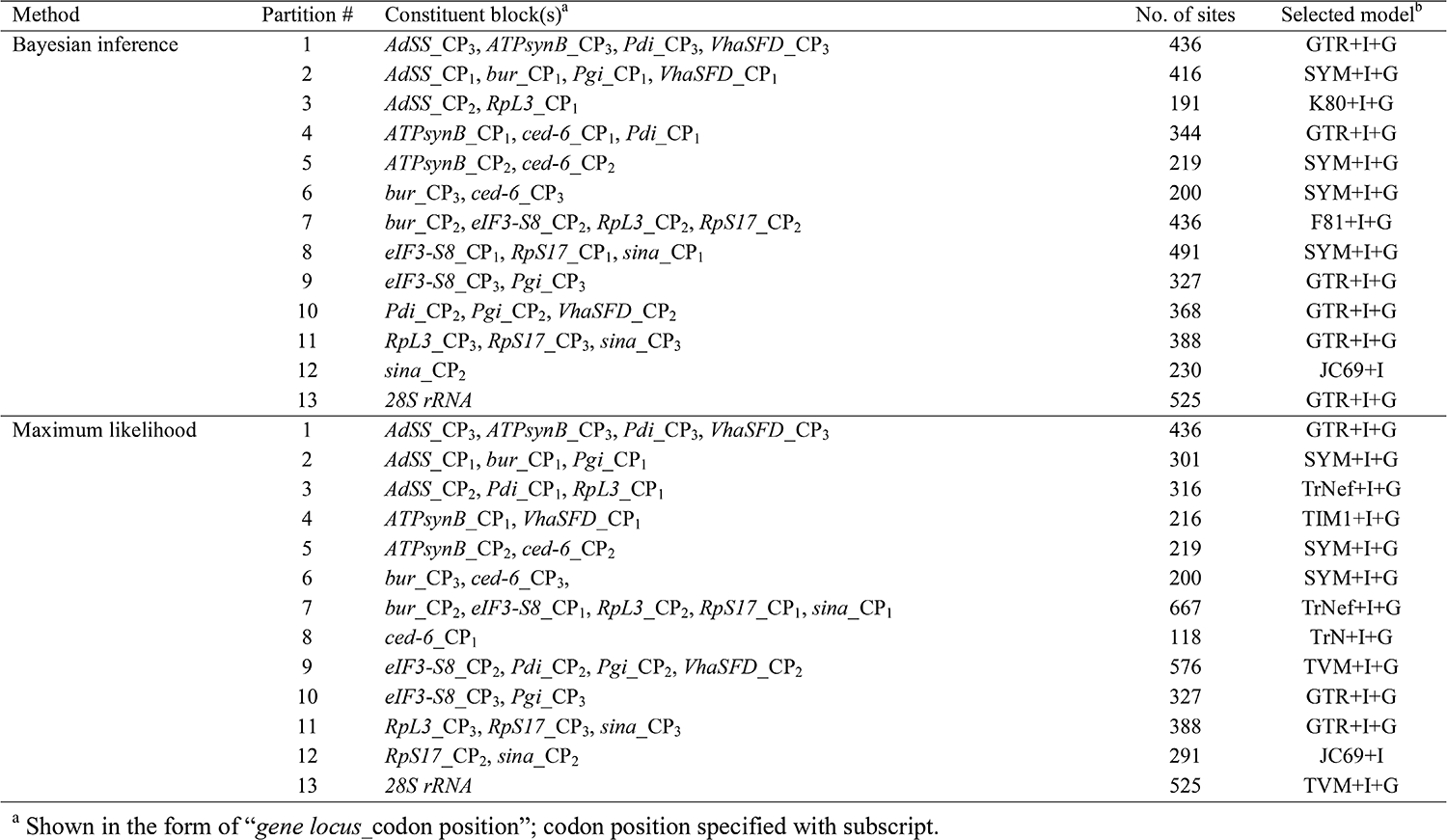
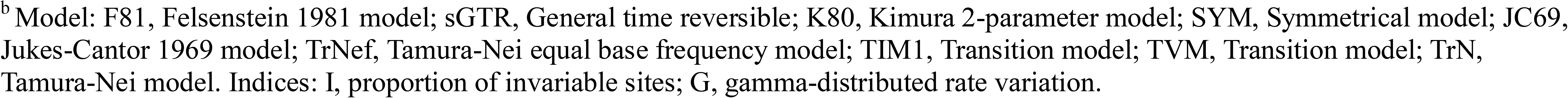
Optimal partitioning strategies and substitution models selected for the concatenated DNA sequences of 12 nuclear gene markers in phylogenetic reconstruction by the Bayesian inference and the maximum likelihood methods.

| Method | Partition # | Constituent block(s)<sup>a</sup> | No. of sites | Selected model<sup>b</sup> |
| --- | --- | --- | --- | --- |
| Bayesian inference | 1 | *AdSS*_CP<sub>3</sub>, *ATPsynB*_CP<sub>3</sub>, *Pdi*_CP<sub>3</sub>, *VhaSFD*_CP<sub>3</sub> | 436 | GTR+I+G |
|  | 2 | *AdSS*_CP<sub>1</sub>, *bur*_CP<sub>1</sub>, *Pgi*_CP<sub>1</sub>, *VhaSFD*_CP<sub>1</sub> | 416 | SYM+I+G |
|  | 3 | *AdSS*_CP<sub>2</sub>, *RpL3*_CP<sub>1</sub> | 191 | K80+I+G |
|  | 4 | *ATPsynB*_CP<sub>1</sub>, *ced-6*_CP<sub>1</sub>, *Pdi*_CP<sub>1</sub> | 344 | GTR+I+G |
|  | 5 | *ATPsynB*_CP<sub>2</sub>, *ced-6*_CP<sub>2</sub> | 219 | SYM+I+G |
|  | 6 | *bur*_CP<sub>3</sub>, *ced-6*_CP<sub>3</sub> | 200 | SYM+I+G |
|  | 7 | *bur*_CP<sub>2</sub>, *eIF3-S8*_CP<sub>2</sub>, *RpL3*_CP<sub>2</sub>, *RpS17*_CP<sub>2</sub> | 436 | F81+I+G |
|  | 8 | *eIF3-S8*_CP<sub>1</sub>, *RpS17*_CP<sub>1</sub>, *sina*_CP<sub>1</sub> | 491 | SYM+I+G |
|  | 9 | *eIF3-S8*_CP<sub>3</sub>, *Pgi*_CP<sub>3</sub> | 327 | GTR+I+G |
|  | 10 | *Pdi*_CP<sub>2</sub>, *Pgi*_CP<sub>2</sub>, *VhaSFD*_CP<sub>2</sub> | 368 | GTR+I+G |
|  | 11 | *RpL3*_CP<sub>3</sub>, *RpS17*_CP<sub>3</sub>, *sina*_CP<sub>3</sub> | 388 | GTR+I+G |
|  | 12 | *sina*_CP<sub>2</sub> | 230 | JC69+I |
|  | 13 | *28S rRNA* | 525 | GTR+I+G |
| Maximum likelihood | 1 | *AdSS*_CP<sub>3</sub>, *ATPsynB*_CP<sub>3</sub>, *Pdi*_CP<sub>3</sub>, *VhaSFD*_CP<sub>3</sub> | 436 | GTR+I+G |
|  | 2 | *AdSS*_CP<sub>1</sub>, *bur*_CP<sub>1</sub>, *Pgi*_CP<sub>1</sub> | 301 | SYM+I+G |
|  | 3 | *AdSS*_CP<sub>2</sub>, *Pdi*_CP<sub>1</sub>, *RpL3*_CP<sub>1</sub> | 316 | TrNef+I+G |
|  | 4 | *ATPsynB*_CP<sub>1</sub>, *VhaSFD*_CP<sub>1</sub> | 216 | TIM1+I+G |
|  | 5 | *ATPsynB*_CP<sub>2</sub>, *ced-6*_CP<sub>2</sub> | 219 | SYM+I+G |
|  | 6 | *bur*_CP<sub>3</sub>, *ced-6*_CP<sub>3</sub>, | 200 | SYM+I+G |
|  | 7 | *bur*_CP<sub>2</sub>, *eIF3-S8*_CP<sub>1</sub>, *RpL3*_CP<sub>2</sub>, *RpS17*_CP<sub>1</sub>, *sina*_CP<sub>1</sub> | 667 | TrNef+I+G |
|  | 8 | *ced-6*_CP<sub>1</sub> | 118 | TrN+I+G |
|  | 9 | *eIF3-S8*_CP<sub>2</sub>, *Pdi*_CP<sub>2</sub>, *Pgi*_CP<sub>2</sub>, *VhaSFD*_CP<sub>2</sub> | 576 | TVM+I+G |
|  | 10 | *eIF3-S8*_CP<sub>3</sub>, *Pgi*_CP<sub>3</sub> | 327 | GTR+I+G |
|  | 11 | *RpL3*_CP<sub>3</sub>, *RpS17*_CP<sub>3</sub>, *sina*_CP<sub>3</sub> | 388 | GTR+I+G |
|  | 12 | *RpS17*_CP<sub>2</sub>, *sina*_CP<sub>2</sub> | 291 | JC69+I |
|  | 13 | *28S rRNA* | 525 | TVM+I+G |

<sup>a</sup> Shown in the form of “*gene locus\_codon position*”; codon position specified with subscript.
<sup>b</sup>Model: F81, Felsenstein 1981 model; sGTR, General time reversible; K80, Kimura 2-parameter model; SYM, Symmetrical model; JC69, Jukes-Cantor 1969 model; TrNef, Tamura-Nei equal base frequency model; TIM1, Transition model; TVM, Transition model; TrN, Tamura-Nei model. Indices: I, proportion of invariable sites; G, gamma-distributed rate variation.

The resulting ML tree rooted with the out-group is shown in Fig. 2. The topology of the BI tree was nearly identical to that of the ML tree, with only minor differences in branching order of the genus *Microdrosophila* Malloch, the subgenus *Siphlodora* of *Drosophila* and a few terminal MOTUs of the *Di. agbo* group (Fig. S2), which have no influence on our conclusions. The three focal taxa, *Dichaetophora*, *Dudaica* and *Mulgravea*, formed a clade with moderate (BP: bootstrap percentage = 67) and strong (PP: posterior probability = 1.00) support in the ML and Bayesian trees, respectively. This clade, hence called the “pan-*Dichaetophora*”, was placed as sister to the moderately supported (BP = 59, PP = 0.74) clade of the *Zygothrica* genus group, which included *Hirtodrosophila*, *Zygothrica* and *Mycodrosophila* in the present study, within the in-group (i.e., the tribe Drosophilini), although this sister relationship was not so strongly supported (BP = 28, PP = 0.63).

**Figure 2.**
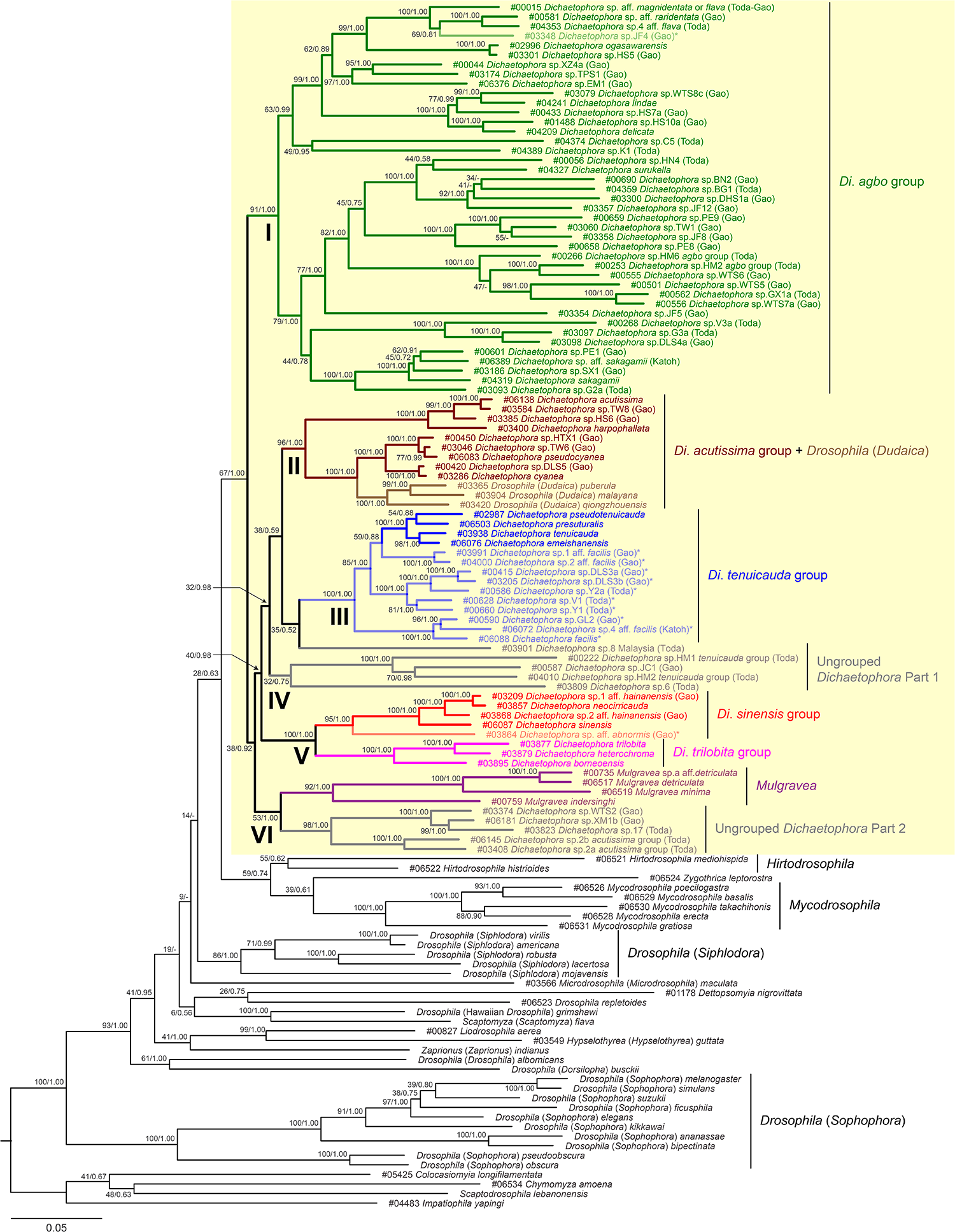
Maximum likelihood tree constructed for the genera *Dichaetophora* and *Mulgravea* and the subgenus *Dudaica* of *Drosophila* as the main target (shaded yellow) in the tribe Drosophilini (in-group), based on DNA sequences of 12 nuclear gene markers. The tree is rooted by out-group rooting (the tribe Colocasiomyini as the out-group). For the three focal taxa, a total of 88 MOTUs (molecular operational taxonomic units) were selected from those inferred from the ABGD analysis for *COI* barcoding (Fig. 1). Supraspecific classification of them based on the morphological diagnosis for each taxon is shown in different colors: the *agbo* (green), *acutissima* (brown), *tenuicauda* (blue), *trilobita* (pink), *sinensis* (red) species groups and ungrouped species (gray) of *Dichaetophora*, *Mulgravea* (purple), and *Dudaica* (light brown). MOTUs only partially concordant with the diagnosis are asterisked and shown in pale color. Support value of ML bootstrap percentage/Bayesian posterior probability (“-” indicating the corresponding branch not realized in the Bayesian inference) is shown for every internal branch. The voucher numbers of specimens used for DNA sequencing in the present study are shown at corresponding terminal branches.

Within the pan-*Dichaetophora*, six clades were recognized. Clade I (BP = 91, PP = 1.00) was comprised of 41 MOTUs, all of which were morphologically classified into the *Di. agbo* group, including five described species, *Di. ogasawarensis*, *Di. lindae*, *Di. delicata* (Nishiharu, 1981), *Di. surukella* (Okada, 1965) and *Di. sakagamii* (Toda, 1989). Clade II (BP = 96; PP = 1.00) consisted of the *Di. acutissima* group and the subgenus *Dudaica*. Within this clade, *Dudaica* was recovered as a monophyletic group (BP = 100, PP = 1.00), but the *acutissima* group was regarded as paraphyletic with respect to *Dudaica*. The *acutissima* group was divided into two solid subclades (BP = 100, PP = 1.00 for both): one including *Di. cyanea* (Okada, 1988) and *Di. pseudocyanea* was placed as sister to *Dudaica* (BP = 100, PP = 1.00), while the other including *Di. acutissima* and *Di. harpophallata* (Hu, Watabe & Toda, 1999) as basal. Clade III (BP = 100, PP = 1.00) corresponded to the *Di. tenuicauda* group. Within this clade, the four known species, *Di. tenuicauda*, *Di. emeishanensis* (Hu & Toda, 1999), *Di. pseudotenuicauda* (Toda, 1983) and *Di. presuturalis* (Hu & Toda, 1999), sharing all the diagnostic characters for this species group, formed a solid cluster (BP = 100, PP = 1.00), but the remaining MOTUs including *Di. facilis* only partially conformed to the diagnosis. The ungrouped MOTU “*Di*. sp.8 Malaysia (Toda)” was placed as sister to Clade III, although the support for this relationship was weak (BP = 35, PP = 0.52). Four other ungrouped MOTUs of *Dichaetophora* formed Clade IV with less support (BP = 32, PP = 0.75). Clade V (BP = 100, PP = 1.00) was comprised of two monophyletic, sister groups, i.e., the *Di. sinensis* group (BP = 95, PP = 1.00) and *Di. trilobita* group (BP = 100, PP = 1.00). The classification of *Di*. *neocirricauda* into the *sinensis* group was corroborated by its morphological congruence with the diagnosis of the *sinensis* group and molecular phylogenetic position, although this species is assigned to the *agbo* group in the current classification (DrosWLD-Species; TaxoDros). Clade VI (BP = 53, PP = 1.00) was also comprised of two sister subclades: one corresponded to the genus *Mulgravea* (BP = 92, PP = 1.00), and the other consisted of five ungrouped *Dichaetophora* MOTUs (BP = 98, PP = 1.00). However, relationships among these six clades of the pan-*Dichaetophora* were less resolved in the present analysis.

## Discussion

Our *COI* barcoding by the ABGD analysis estimated 222 MOTUs among a total of 1,070 specimens of the three taxa, the genera *Dichaetophora* and *Mulgravea* and the subgenus *Dudaica* of *Drosophila*, on which the present study focused. Most of these MOTUs corresponded to morpho-species first distinguished. The 34 known species included in the analysis were all recognized as individual MOTUs. Among them, *Di. delicata*, which had been synonymized, along with *Di. pleurostriata* (Singh & Gupta, 1981), with *Di. lindae* by Okada (1984a), was included as a distinct MOTU (Figs 1 and 2, Fig. S1): thus, its status as a valid species was strongly suggested by both morphological and molecular evidence. The status for these three species once synonymized should be reconsidered in connection with other related MOTUs inferred from the present study. On the other hand, some morpho-species were divided into multiple MOTUs: for example, “*Di*. sp.G2 (Toda)” into sp.G2a and sp.G2b MOTUs, “*Di*. sp.G3 (Toda)” into sp.G3a and sp.G3b MOTUs, and “*Di*. sp.WTS8 (Gao)” into sp.WTS8a–d MOTUs (Fig. 1, Fig. S1). These MOTUs may include cryptic, sibling species, subspecies, or intraspecific genetic varieties. On the contrary, two morpho-species, “*Di.* sp.DLS3a (Gao)” and “*Di.* sp.DLS3b (Gao)” which are distinctly different from each other in the morphology of aedeagus and aedeagal apodeme (Fig. 1H), were combined into a single MOTU (Fig. 1, Fig. S1). To determine boundaries of valid species, we need to re-examine the morphology of suggested MOTUs in more detail and to incorporate data from biogeography and ecology into the species delimitation. Leaving such a work for the final delimitation to Part II of this serial study, we tentatively classified the suggested MOTUs (222 in total) into supraspecific categories (genus, subgenus and/or species group) under the current classification framework of the focal taxa: 128 of the *agbo* group, 39 of the *tenuicauda* group, 18 of the *acutissima* group, 6 of the *sinensis* group, 6 of the *trilobita* group, and 12 ungrouped species of *Dichaetophora*, 5 of *Mulgravea*, and 8 of *Dudaica* (Table 4).

**Table 4.** The numbers of known species and MOTUs (molecular operational taxonomic unit) estimated by the ABGD analysis for *COI* barcoding, separately enumerated for each species group and ungrouped category of *Dichaetophora*, the genus *Mulgravea*, and the subgenus *Dudaica* of *Drosophila*.

| Genus (subgenus) and species group | Number of known species <sup>a</sup> | MOTUs<sup>b</sup> estimated by ABGD / subjected to phylogenetic reconstruction |  |  |
| --- | --- | --- | --- | --- |
|  |  | Known species | Newly recognized | Total |
| *Dichaetophora agbo* group | 43 | 6/5 | 122/36 | 128/41 |
| *Dichaetophora tenuicauda* group | 10 | 6/5 | 33/8 | 39/13 |
| *Dichaetophora acutissima* group | 4 | 4/4 | 14/5 | 18/9 |
| *Dichaetophora sinensis* group | 4 | 2/2 | 4/3 | 6/5 |
| *Dichaetophora trilobita* group | 6 | 6/3 | 0/0 | 6/3 |
| Ungrouped *Dichaetophora* | 0 | 0/0 | 12/10 | 12/10 |
| *Mulgravea* | 14 | 3/3 | 2/1 | 5/4 |
| *Drosophila* (*Dudaica*) | 8 | 7/3 | 1/0 | 8/3 |
| **Total** | **89** | **34/25** | **188/63** | **222/88** |

<sup>a</sup> Based on the current classification of DrosWLD-Species and TaxoDros.
<sup>b</sup> Classified into supraspecific taxa (genus, subgenus and/or species group), based on the morphological diagnosis for each taxon.

Of these MOTUs, 88 (41 of the *agbo* group, 13 of the *tenuicauda* group, 9 of the *acutissima* group, 5 of the *sinensis* group, 3 of the *trilobita* group, and 10 ungrouped species of *Dichaetophora*, 4 of *Mulgravea*, and 3 of *Dudaica*) were selected for the phylogenetic reconstruction based on the 12 nuclear gene markers. In the resulting tree, the three focal taxa (*Dichaetophora*, *Mulgravea* and *Dudaica*) formed a clade, which we provisionally called the pan-*Dichaetophora*. Within this large clade, *Mulgravea* and *Dudaica* were recovered as monophyletic groups, but *Dichaetophora* was paraphyletic in respect to the former two taxa. Furthermore, four of the five species groups of *Dichaetophora* were recovered as strongly supported monophyletic groups, but the *acutissima* group was paraphyletic in respect to *Dudaica*. Of the four monophyletic species groups, the *sinensis* and *trilobita* groups were sister to each other. In addition, two solid clades were recognized among the ungrouped MOTUs of *Dichaetophora*. One was comprised of five MOTUs of Part 2 and sister to *Mulgravea*, and the other was of three MOTUs tentatively classified into Part 1. Thus, the present study has uncovered some problems in the taxonomy of the pan-*Dichaetophora*. For instance, (i) are *Mulgravea* and *Dudaica* to be synonymized with *Dichaetophora*, or is the pan-*Dichaetophora* to be divided into a few genera; (ii) are the *sinensis* and *trilobita* groups to be combined into a species group; and (iii) are any species subgroups and/or complexes to be established within species groups? To solve these problems, it would be effective to incorporate all known and putatively new species (i.e., MOTUs inferred from the present study) into a grafting phylogenetic analysis with morphological characters (e.g. Fu *et al.*, 2016). Based on the topology of resulting tree and the character mapping on it, the clade-based classification system will be established for the pan-*Dichaetophora*: each supraspecific taxon is to be monophyletic and defined by synapomorphies as the diagnosis, and all component species are explicitly assigned in the hierarchical system of such supraspecific taxa. This will be challenged in Part II of this serial work.

## Acknowledgements

We thank Administrations of the Xishuangbanna, Gaoligongshan, Jianfengling and Taibaishan National Nature Reserves, the Xishuangbanna Tropical Botanical Garden, CAS, the Fushan Botanical Garden, and the Forestry Bureau of Xizang Autonomous Region for administrative supports during field survey in each Garden/Reserve. We also thank Dr. Shun-Chern Tsaur, Dr. Chau-Ti Ting, Dr. Fu-Guo Robert Liu, Mr. Zhi-Wei Chang, Dr. Hong-Wei Chen, Dr. Xi-Peng Chen, Mr. Zhao Fu, Ms. Nan-Nan Li for their help in field works in China; Dr. Shin-ichi Tanabe, Dr. Maklarin B. Lakim, Dr. Rimi Repin and Dr. Maryati Bte Mohamed for their help in field works in Sabah, Malaysia, under research permissions (UPE:40/200/19 SJ. 732 and UPE: 40/200/19 SJ.1194 and 1195) of Economic Planning Unit of Malaysian Government; Dr. Lucy Chong, Dr. Kohei T. Takano, Dr. Takao Itioka and Dr. Tohru Nakashizuka for their help with field work in the Lambir Hills National Park, Sarawak, Malaysia, in accordance with the Memorandums of Understanding signed between the Sarawak Forestry Corporation (SFC, Kuching, Malaysia) and the Japan Research Consortium for Tropical Forests in Sarawak (JRCTS, Sendai, Japan) in November 2005, and under Sarawak Forestry Department Permission to Conduct Research on Biological Resources – Permit No. NCCD.907.4.4(J|d.7)-109 and Park Permit No. 50/2012; Dr. Sri Hartini and other staff members of Zoology Division, Research Center for Biology-LIPI, Indonesia, for their help with field work in Java and Sumatra under research permissions 5816/SU/KS/2004 and 6967/SU/KS/2004; Dr. Bui Tuan Viet for his help with field work in Vietnam; Prof. Masayoshi Watada for his help in field works in Ehime, Japan; and Dr. Artyom Kopp and Dr. Toru Katoh for discussion about phylogenetic reconstruction. This work was supported by NSFC (Nos 31160429, 32060112 and 31760617), the fund of the Ministry of Science and Technology of China (Nos 2011FY120200 and 2012FY110800), JSPS KAKENHI Grants Numbers 15255006, 21570085, 23405004, 24370033 and 25257416, Creative Basic Research 09NP1501 and the 21st Century Center of Excellence Program (E-01) of the Ministry of Education, Culture, Sports, Science and Technology, Japan.

## Supplementary information

**Table S1.** Specimens of the genera *Dichaetophora* and *Mulgravea* and the subgenus *Dudaica* of *Drosophila* used for *COI* barcoding.

**Table S2.** Species/MOTUs and DNA sequences (newly determined ones: MT662140–MT663148) employed in the phylogenetic reconstruction. A hyphen (“-”) is used to indicate a case of missing data.

**Figure S1.** Neighbor-joining tree (original) built with 1,070 *COI* sequences of the genera *Dichaetophora* and *Mulgravea* and the subgenus *Dudaica* of *Drosophila*. Voucher specimen number (or GenBank accession number) is shown for each sequence. MOTUs inferred from the ABGD analysis is shown, with indication of supraspecific classification in different colors: the *agbo* (green), *acutissima* (brown), *tenuicauda* (blue), *trilobita* (pink), *sinensis* (red) species groups and ungrouped species (gray) of *Dichaetophora*, *Mulgravea* (purple), and *Dudaica* (light brown). MOTUs only partially concordant with the diagnosis are asterisked and those having female specimens only are indicated by dagger (†), and both are shown in pale color. MOTUs (voucher or Genbank accession numbers of the selected specimens underlined) subjected to the phylogenetic reconstruction are shown with thick branch(es). Bootstrap percentages more than 50% are shown beside nodes.

**Figure S2.** Bayesian Inference tree constructed based on DNA sequences of 12 nuclear gene markers. Support value of Bayesian posterior probability is shown for every internal branch. The dashed lines indicate branches resulting only from the ML analysis. See the legend of Fig. 2 for more explanation.

