## Supplemental Figure 1 for "Revision of the genus *Dichaetophora* Duda (Diptera: Drosophilidae), part I: DNA barcoding and molecular phylogenetic reconstruction"

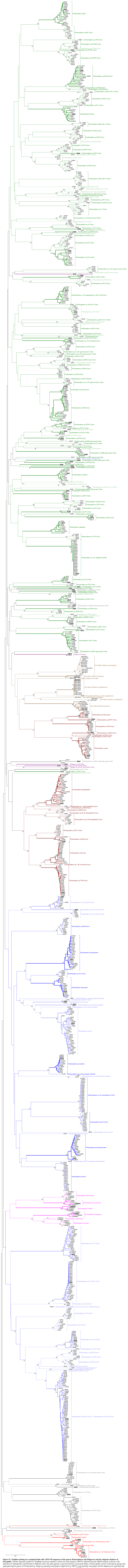

**Figure S1. Neighbor-joining tree (original) built with 1,070 *COI* sequences of the genera *Dichaetophora* and *Mulgravea* and the subgenus *Dudaica* of *Drosophila*.** Voucher specimen number (or GenBank accession number) is shown for each sequence. MOTUs inferred from the ABGD analysis is shown, with indication of supraspecific classification in different colors: the *agbo* (green), *acutissima* (brown), *tenuicauda* (blue), *trilobita* (pink), *sinensis* (red) species groups and ungrouped species (gray) of *Dichaetophora*, *Mulgravea* (purple), and *Dudaica* (light brown). MOTUs only partially concordant with the diagnosis are asterisked and those having female specimens only are indicated with dagger (†), and both are shown in pale color. MOTUs (voucher or Genbank accession numbers of the selected specimens underlined) subjected to the phylogeny reconstruction are shown with thick branch(es). Bootstrap percentages more than 50% are shown beside nodes.
