## Supplementary figures and images for "Revision of the genus *Dichaetophora* Duda (Diptera: Drosophilidae), part I: DNA barcoding and molecular phylogenetic reconstruction"

### Supplemental Figure 2

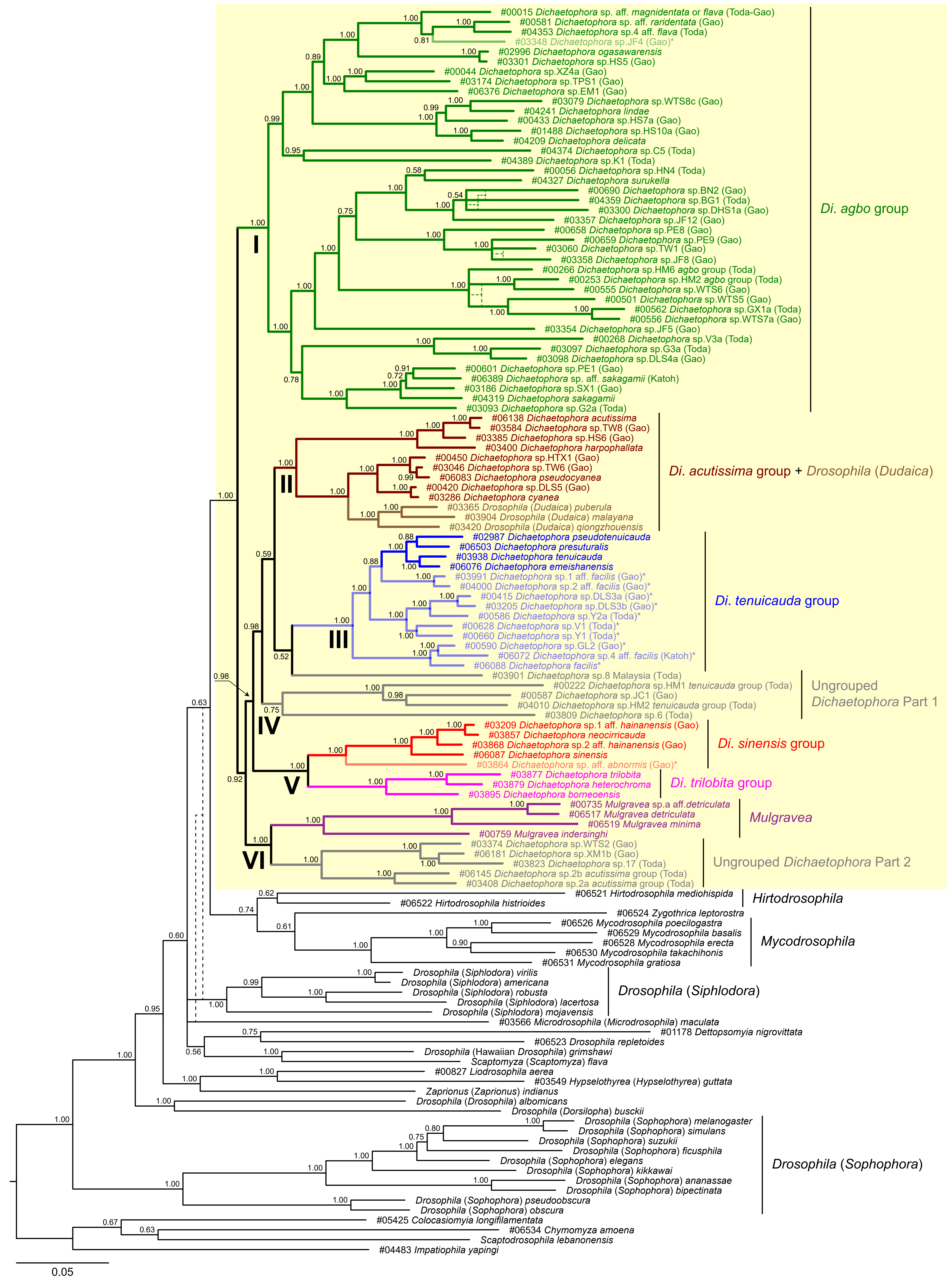
